# Advanced processing and analysis of conventional confocal microscopy generated scanning FCS data

**DOI:** 10.1101/163766

**Authors:** Dominic Waithe, Falk Schneider, Jakub Chojnacki, Mathias Clausen, Dilip Shrestha, Jorge Bernardino de la Serna, Christian Eggeling

## Abstract

Scanning Fluorescence Correlation Spectroscopy (scanning FCS) is a variant of conventional point FCS that allows molecular diffusion at multiple locations to be measured simultaneously. It enables disclosure of potential spatial heterogeneity in molecular diffusion dynamics and also the acquisition of a large amount of FCS data at the same time, providing large statistical accuracy. Here, we optimize the processing and analysis of these large-scale acquired sets of FCS data. On one hand we present FoCuS-scan, scanning FCS software that provides an end-to-end solution for processing and analysing scanning data acquired on commercial turnkey confocal systems. On the other hand, we provide a thorough characterisation of large-scale scanning FCS data over its intended time-scales and applications and propose a unique solution for the bias and variance observed when studying slowly diffusing species. Our manuscript enables researchers to straightforwardly utilise scanning FCS as a powerful technique for measuring diffusion across a broad range of physiologically relevant length scales without specialised hardware or expensive software.

## Introduction

FCS (Fluorescence Correlation Spectroscopy) is a long established technique which allows the diffusion characteristics of fluorescently labelled molecules to be measured (Magde, Elson et al. 1972, Ehrenberg and Rigler 1974). In FCS the molecular mobility is accessed by determining the time it takes the molecules under study to traverse through the observation spot of an optical microscope. Knowledge of both the average transit time and the size of the observation spot allows a straightforward determination of respective molecular diffusion coefficients as well as anomalies in diffusion. Through the advent of confocal microscopy, FCS has become a powerful tool in various fields of application (Rigler and Widengren 1990). An important topic is the use of FCS for determining the diffusion dynamics of molecules and proteins in the membrane of cells and lipid bilayers (Fahey, Koppel et al. 1977, Widengren and Rigler 1998, Schwille, Korlach et al. 1999). For FCS to fundamentally establish the dynamics of proteins in living membranes however, spatial information as well as high statistical accuracy through large data sets are required, due to the heterogeneity and dynamic nature of the membrane under study (Lingwood and Simons 2010, de la Serna, Schütz et al. 2016, Sezgin, Levental et al. 2017). Conventional (point) FCS employs a single measurement spot at a time only. In this paradigm a laser is focused on a single spot in the sample and the emission light is recorded at this location throughout the duration of the experiment. As a consequence spatial information and sufficient statistical power, perquisites of studying complex heterogeneous organisms, can only be achieved through subsequent measurements. With measurement times of seconds to tens of seconds a sequential approach like point FCS often results in averaging of possible diffusion dynamics and so obscuring the vital diffusion heterogeneities that are of interest.

The most obvious way of attaining spatial information, as well as large data sets, is through the recording of FCS data in multiple observation spots simultaneously. Through simultaneous measurement, heterogeneities present between locations become much more obvious due to the high-degree of temporal synchronisation allowing accurate comparison. Simultaneous measurements can be achieved through camera-based (image correlation spectroscopy, ICS) (Petersen, Höddelius et al. 1993) or multi-spot approaches (Brinkmeier, Dörre et al. 1999, Colyer, Scalia et al. 2010, Papadopoulos, Krmpot et al. 2015). While the spatial awareness of these approaches have allowed the study of a wide range of cellular processes, especially through there further development (e.g. (Digman, Sengupta et al. 2005, Hebert, Costantino et al. 2005, Sankaran, Manna et al. 2009), the general applicability of each is hampered either by the low frame rate of the detectors, limiting e.g. the use of ICS for slow diffusion processes only, or by a complex setup. Scanning FCS is a variant of the conventional point FCS technique and allows diffusion at multiple locations to be measured simultaneously on conventional light microscopy equipment (Ruan, Cheng et al. 2004, Ries, Chiantia et al. 2009, Honigmann, Mueller et al. 2014). In scanning FCS, the laser illumination spot is scanned across a sample in a line (or circle) continuously and the light emitted from each spatial location recorded in sequence. The intensity time-series at each point is then correlated and parameterised through fitting of derived equations which link diffusion dynamics with the relaxation of the correlation function. The yielded diffusion data can then be used to establish spatial relationships within the sample. The compromise of this technique is still a reduced temporal resolution, as the illumination and detection spot can only be in one location at one time when compared to point FCS, i.e. the temporal resolution is ultimately given by the scanning frequency and the signal-to-noise ratio. Still, compared with other ICS-based approaches, scanning FCS offers a good compromise for producing both spatial and high frequency temporal information with the added benefit that it can be performed on conventional laser scanning microscope equipment.

Noise and its likely impact on a measurement is an important consideration when performing correlation, and several attempts to characterize noise in FCS data analytically have been made in the past. Koppel first showed that for experiments of a long duration that the main basis for achieving a good signal-to-noise ratio was to have a strong signal per molecule when compared to background noise like shot noise (Koppel 1974). Subsequent attempts to understand the statistical accuracy of correlation have highlighted the importance of the duration of an experiment and also the artifacts of having finite measurement duration (Schätzel, Drewel et al. 1988, Saffarian and Elson 2003). Knowledge of the relative levels of error across a correlation function have been shown to assist in the accuracy of curve fitting and increasing the accuracy of diffusion coefficient calculation (Wohland, Rigler et al. 2001). Scanning FCS may give further insights into such issues as it allows acquisition of large-scale FCS data sets simultaneously.

Here, we aim to optimize both processing and analysis for large-scale scanning FCS data by on the one hand presenting the novel FoCuS-scan software and secondly by providing a thorough characterization of scanning FCS in a typical biological user-case. FoCuS-scan software provides an end-to-end solution for processing and analysing scanning data produced using commercial turnkey systems and expands existing software for processing scanning FCS samples (Rossow, Sasaki et al. 2010, Müller, Schwille et al. 2014) with respect to ease-of-use, flexibility and analysis of large data batches. FoCuS-scan software contains tools that allow common photobleaching artefacts to be compensated for, as well as tools that allow cropping to be applied to samples and innovative visualization techniques. Furthermore, the FoCuS-scan software also utilises advanced fitting algorithms, which accompany the data processing and allow large and complex datasets to be efficiently analysed. Using this software, we in this study further characterize scanning FCS for use in investigating slow moving molecular species on individual cells (1.0 μm^2^/s to 0.05 μm^2^/s, much slower than has been previously characterized in FCS statistical analysis). Due to the power of FoCuS-scan and the applicability to turn-key confocal systems it is an essential tool for any bioscience researcher interested in studying cellular dynamics.

Toward our goal of optimizing processing and analysis of scanning FCS data we characterize simulations generated to model typical biological acquisition experiments. These simulations highlight the limitations of experiments where there is a limited number of molecules and also a need to maximize the number of acquisitions possible within a period. As a consequence we have simulated acquisition times of varying length and also traces that model photobleaching artifacts. We observe that in a typical biological user-case the population of transit times significantly broaden with decreasing theoretical diffusion rate especially at slow diffusion rates in combination with a limited acquisition time. This phenomenon is systematic of a limited convergence of the underlying molecular motion being analyzed. Within the results section we characterize the statistical accuracy of using scanning FCS in this situation and propose solutions that can improve the resolution of the technique for resolving slower moving species as well as powerful means to visualize and understand the data.

## Methods

### Acquisition and Simulation

#### Cell culturing and preparation for microscope

Jurkat T-cells were cultured in RPMI-1640 (Sigma Aldrich, UK) media supplemented with 10% FBS (Sigma Aldrich), 2 mM L-Glutamine (Sigma Aldrich), 100 U/mL Penicillin (Sigma Aldrich), 0.1 mg/mL Streptomycin (Sigma Aldrich) and 10 mM HEPES pH 7.4 (Sigma Aldrich). 1 million cells were spun down for 5 minutes at 2000 rpm and washed with 1 mL of L15 medium (Life Technologies). After spinning down again the cells were labelled by resuspending in L15 medium containing 0.4 ug/mL Atto647N-DPPE (AttoTec). The cells were labelled at 37 C shaking at 300 rpm for 15 minutes. After washing with L15, the cells were resuspended and kept in L15 for not longer than 1 hour on room temperature. Measurements were performed in 8-well glass-bottom chambers (Ibidi). Prior to the measurements the glass was coated with PLL using a 0.01 % PLL-solution (Poly-L-Lysine) (Sigma Aldrich) for 1 hour at room temperature and washed three times with L15.

#### Scanning FCS experimental acquisition

The scanning FCS measurements were performed using a customized Abberior Instruments microscope (Galiani, Waithe et al. 2016). The microscope was controlled by Abberior’s Imspector software. Scanning FCS measurements were acquired in xt mode using an orbital scan with a pixel dwell time of 10 µs and a scanning frequency of 2630 Hz. The pixel size was typically in the range of 80-150 nm and the diameter (full-width-at-half-maximum, FWHM) of the observation spot (point-spread-function, PSF) was taken to be that of a conventional confocal microscope (250 nm). The fluorescence was excited using a 640 nm pulsed diode laser (PicoQuant) with 10 µW total excitation power at the objective’s back aperture. The detector used was an Excelitas APD (SPCM-AQRH-13).

#### Scanning FCS experimental processing

For a scanning FCS measurement the fluorescence intensity of the sample across a laser line was systematically collected for the duration an experiment (Fig. 1A). The laser focus was moved over the specimen, recording at each of M positions, before repeating the cycle N times. The duration spent scanning across a pixel location (e.g. M=0) is known as the dwell time, whereas the time taken to repeat one of N cycles is denoted the line time. On most current commercial turnkey systems the laser can only be configured to scan along a line on the sample and so there is delay whilst the laser focus is moved back to the origin. With the correct equipment however (e.g. in Fig. 1A-B), it is possible to scan a circle, in which case the summed dwell time for all M locations (ΣM) and the line time will be the same. The length of the line scanned was typically around 5 μm with often M=64 locations specified. Experiments were varied in duration, but between 10-60 s was a practical range depending on the mobility of the species being studied. An intensity carpet, which contains the intensity measurements made at each spatial location and time point is the direct product of a scanning FCS measurement. Intensity carpets were exported from the microscope and then imported into the FoCuS-scan software for correlation and analysis. The FoCuS-scan software accepts file types from the major turn-key system providers such as Leica (.lif), Zeiss (.lsm) or Abberior Instruments (.msr). Nevertheless, using FIJI any other data type can be easily converted to .OME-TIFF which allows to read files from practically every microscopy set-up.

**Figure 1:**
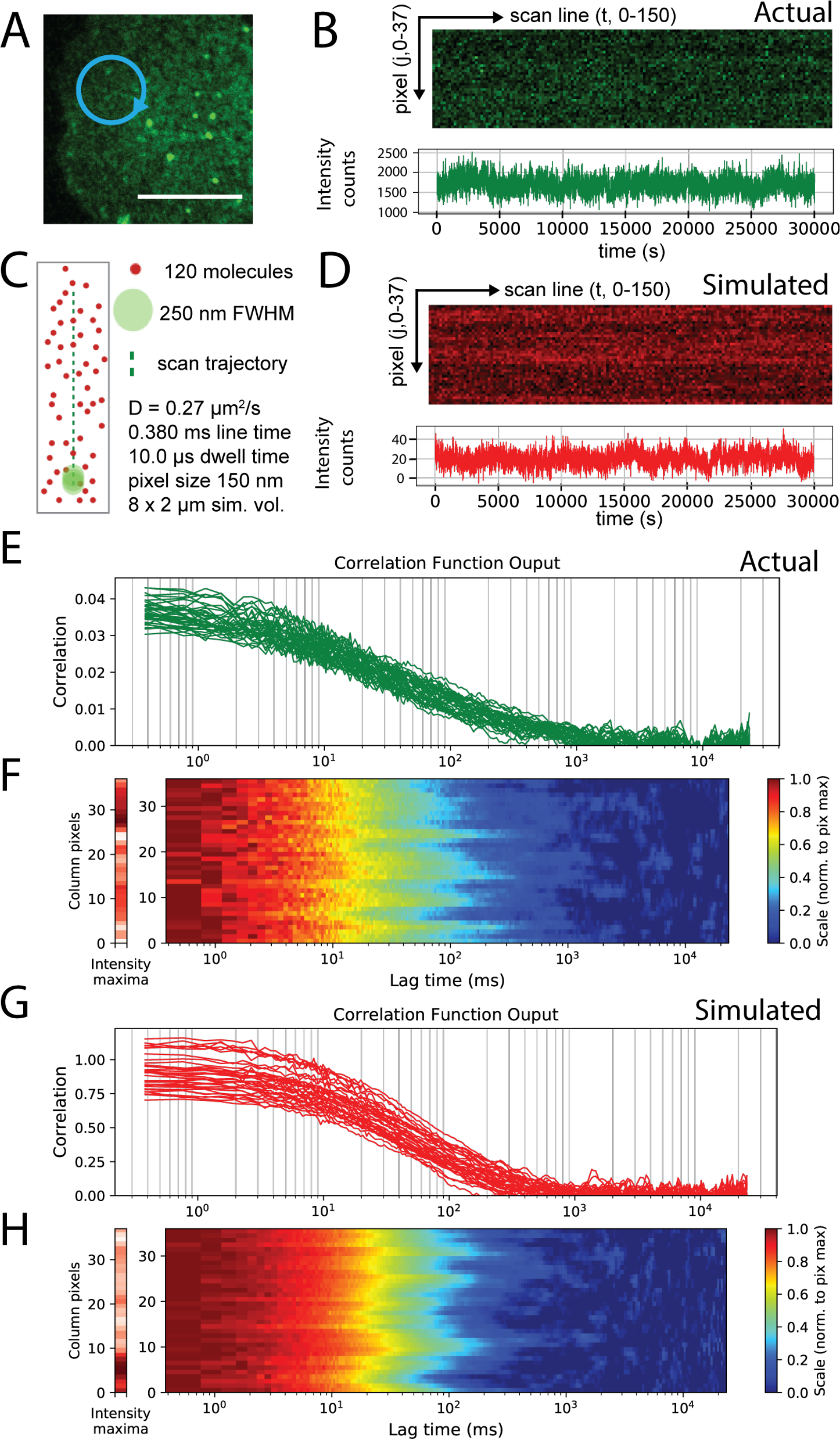
Simulations closely match experimental data. Experimental and simulated scanning FCS data visualized as time-series, correlation plots and correlation carpets**. A**) 2-dimensional scanning confocal monograph of Jurkat T cell surface labelled with Atto647N-DPPE, 10 μm scale (white bar) and representative elliptical scan line (blue circle). **B**) Intensity time-series from the live-cell experiment, (**upper panel**) for all pixels (**y-axis**) and first 150 scan lines (**x-axis**) and (**lower panel**) intensity integrated over all pixels against time. **C**) Schematic representing the molecular simulation and scan line with labeled diffusing molecules (red dots), observation spot (green dot, diameter/FWHM = 250 nm), scan line (green line), and scanning parameters as given. **D**) Intensity time-series from simulated experiment, (**upper panel**) for all pixels (**y-axis**) and first 150 scan lines (**x-axis**) (intensity carpet) and (**lower panel**) intensity integrated over all pixels against time. (**E-H**) Correlation curves yielded from individual pixels positioned along scanning orbit as individual plots (**E** and **G**) or represented as a normalized carpet (**F** and **H**, y-axes: spatial pixel number along scanned line; x-axes: correlation lag times τ left graph: time-integrated intensity for each pixel from 0 (white) to maximum (dark red); red graph: color bar coding of normalized correlation data), for the live cell experiment (**E** and **F**) and the simulated experiment (**G** and **H**).

#### Scanning FCS simulated modelling and acquisition

To explore the parameter space of the scanning FCS technique, we developed simulations that would allow us to model a variety of diffusion modes and acquisition settings. Each simulation modelled a rectangular region (2 by 8 μm in size) on which molecules diffused by 2-dimensional stochastic Brownian motion (Fig. 1C). Diffusion rates between 1.00 and 0.05 μm^2^/s were simulated for durations of between 3000 and 30000 ms. Between 120 and 240 molecules were simulated with a time-step of between 0.002 and 0.0010 ms for each movement. For the simulated acquisition, a Gaussian PSF with a 250 nm FWHM was generated and positioned at different locations along the theoretical scan line with the location and integration time being synchronised with the time-point and scan frequency. Thus it was possible to measure the particles in the same way as in an actual scanning FCS experiment with dwell times and line times which matched the real experiments. For the simulations the scan-line was set to either 5550 or 5812.6 nm in length, with either 37 or 64 points along the line placed at 150 or 90.82 nm intervals respectively. During simulation, when molecules reach the boundary they were wrapped round to the opposite side. The output time-series of one of the simulations is shown in Fig. 1D. We noticed no artefacts with this method and found the support to be of sufficient size to avoid strange effects that might originate from a small simulation area. Shot-noise was simulated by the addition of Gaussian noise (with standard deviation = 1.0) to the data generated at each spatial location of the simulation. The simulations used for this study were written in python and are available as a series of ipython notebooks stored in the Github repository: https://github.com/dwaithe/nanosimpy/simulations

### Software Implementation and algorithms

#### Software Engineering

The FoCuS-scan software is written using the python scripting language and is compiled into an application that can run independently in the Windows, OS X and Linux operating systems. The interface is designed using PyQT library and the bulk of the visualization is performed using the matplotlib visualization library. The fitting of diffusion functions is performed using the lmfit library (Newville, Nelson et al. 2016). Binary file readers for Leica file format (.lif) and the Abberior Imspector software (.msr) have been created and the file readers for the Zeiss (.lsm) and OME-Tiff (.tiff, .tif) files were produced using tifffile python library. The raw python code used for FoCuS-scan and also the compiled binary files are available through: https://github.com/dwaithe/FCS_scanning_correlator

#### Data Pre-processing

Intensity carpets yielded from an experiment or a simulation that are imported into FoCuS-scan are processed in a number of different independent ways. To generate an intensity time-series visualization, the intensity carpets are integrated across all spatial locations to yield an intensity plot over time (Figure 1B):

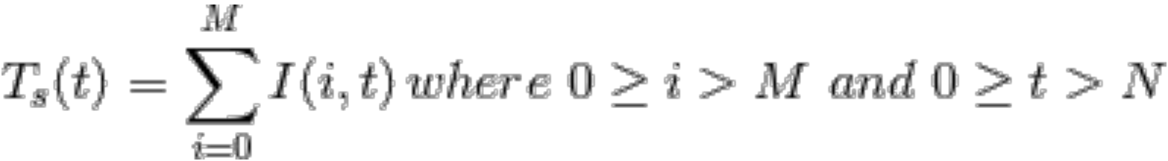

where M is the number of spatial pixels locations in the scan and N is the number of scan line passes. Furthermore, several optional techniques are also possible which can be used to pre-process the data before commitment through correlation. It is possible to crop the data, either in terms of sub-duration or spatially in terms of pixels measured. This cropping facility is included because sometimes only a subset of the data are required for an experiment or it is desirable to exclude artefacts present during certain time spans of the acquisition or certain pixels:

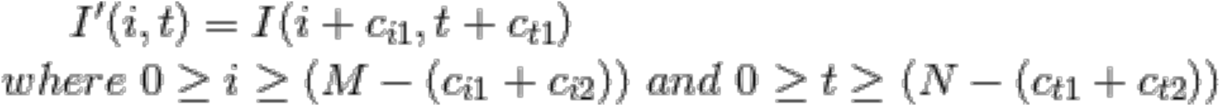

where c_i1_ is the lower crop pixel location and c_i2_ is the greater crop pixel location of the scan line (c_i1_ <c_i2_). Temporally ct1 represents the start time-point from which to start correlation and c_t2_ represents the last time-point from which to correlate (c_t1_ < c_t2_). Within the software interface of FoCuS-scan it is also possible to split the intensity time-series up into multiple sections temporally by setting a desired interval. Processing this step will result in multiple I’ representing differently cropped sections of the input I.

Another optional pre-processing step is to perform spatial binning on the input intensity carpet. This pre-processing step can be applied to reduce the impact of noise from the specimen through the application of a sliding window to the input pixels.

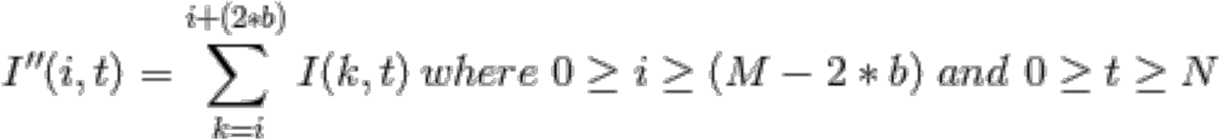

where I’’ is the spatially binned carpet and b is the size of the margin used for the binning.

#### Correlation

Correlation analysis is as signal-processing technique used to determine statistically the time scale that two signals resemble one another. In terms of Fluorescence Correlation Spectroscopy, auto-correlation (G_AC_) represents a signal correlated with itself, the self-similarity over different time-scales, whereas in cross-correlation (G_cc_) two signals are compared from different image channels. Because the signal fluctuations are related to the fluctuations caused by the fluorophores moving in and out of the confocal detection volume, the self-similarity detected within these methods is directly related to the rate of the diffusion of the detected species. Primarily for the software, correlation is performed in time only, but on each pixel j (of the M spatial locations) of the raw intensity data I or on the pre-processed data I’ or I’’ from a specific channel. The auto-correlation carpet is defined as follows:

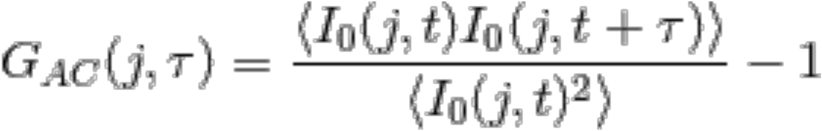

where G_AC_ is the normalised auto-correlation function and I_0_ represents the intensity time-series I for a specific channel (0, or 1), t is the time-point of the time-series, τ is the lag time and j is the scan-line pixel of acquisition. The normalised cross-correlation function GCC represents the correlation calculation for two independent channels I_0_ and I_1_:

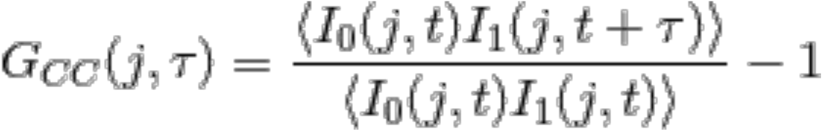

I_0_ and I_1_ would typically represent imaging data acquired at different acquisition wavelengths. The core of the correlation algorithm utilises a fast implementation of a multiple-tau algorithm (Müller 2012). The multiple-tau algorithm was developed more than 30 years ago to produce a method that could efficiently correlate a wide range of delay times using a semi-logarithmic scheme (Schätzel 1985). In FoCuS-scan this multiple-tau algorithm is seamlessly integrated with a single parameter ‘m’ (or quality) that defines the number of points calculated at each level of the logarithmic scheme. Please refer to the supplementary methods for more details regarding parameter selection.

The output of the correlation is displayed as both a correlation trace (Fig. 1C) or as a part of a correlation carpet (Fig. 1D), with the colour of the carpet representing the correlation function output at each correlation lag time. Using the correlation carpet it is more straight-forward to visualize differences between neighbouring pixel locations. These and additional features for processing and visualization can be used according to the user-manual in the Supplementary materials section.

#### Photobleaching correction

FoCuS-scan has two methods for photobleaching correction included. The first method, a common method for correcting for photobleaching, is to correct the decay in the fluorescence time-series T_s_ directly, by fitting with a mono-exponential function (Ries, Chiantia et al. 2009)

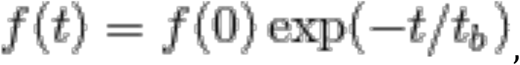

where f(0) is initialised as T_s_(0) and t_b_ denotes the average bleaching time taken for the initially observed fluorescence intensity value to decrease by a factor of 1/e and is extracted by fitting T_s_. Upon determination of f(t) the intensity trace is corrected according to

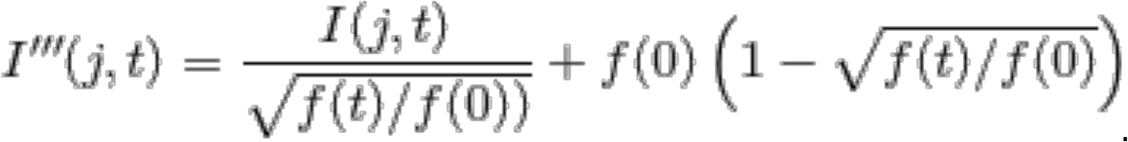

where I(j,t) is the original intensity trace (can be substituted for I’ or I’’) and I’’’(t) is the corrected output. The corrected pixels are then correlated using the standard correlation methodology.

A second method for correction of photobleaching artefacts is to correlate sub-intervals of an input time-series, through local average adaptive correction (Widengren and Rigler 1998, Wachsmuth, Weidemann et al. 2003, Wachsmuth, Conrad et al. 2015). Photobleaching affects correlation especially for lower frequencies, which in turn affects the normalisation of the curve and therefore perturbs the transit time calculation. If rather than compensating for this directly, as in the previous method, it is also possible to avoid the issue by not correlating at long lag times. Because the affect of the photobleaching is predominantly mediated through the longer frequencies it can be circumvented by not correlating up to those times. Because we are no longer correlating long frequencies we can correlate the input time-series at multiple points and improve the statistics of the method by averaging the correlation function resulting from each section. In FoCuS-scan we provide a means to do this and implement it seamlessly as an alternative to the conventional correlation:

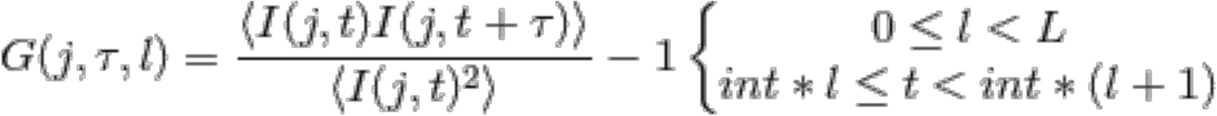

where L represents the number of intervals to divide up our intensity time-series, and *int* is the interval duration (i.e. total duration of time series divided by L). G(j,τ,l) is our correlation function matrix which contains each of the correlation functions. Once the correlation function matrix has been filled, one output correlation function G’ is created which is an average of each of the L interval correlation functions at each lag time and each spatial position:

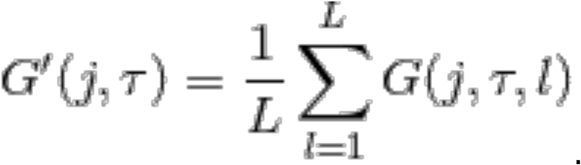

Once correlated this data is exported as normal. The difficulty with this technique is to choose a significantly long interval time to allow the full correlation function decay for a particular species to take shape (convergence of the correlation function) whilst balancing the need to shorten it in favour of reducing the contribution of artefacts resulting from photobleaching of less mobile, or immobile species. FoCuS-scan includes an interface that makes this process straight-forward and reproducible (See supplementary materials for more information).

#### Fitting

Ultimately auto- and cross-correlation functions are calculated to establish the diffusion rates of the species that are responsible for generating the observed functions. Once the correlation function G(j, τ) of an intensity time-series I(j,t) has been calculated we want to determine the characteristic transit time of the observed molecules through the observation spot along with other parameters which describe this function in terms of the underlying physical processes which created it. We do this independently for each individual curve of the carpet rather than considering the whole ensemble and so for simplicity refer to each individual correlated functions as G(τ). Derivations which link G(τ) to the kinetics of fluctuations for 2- and 3-dimensional processes and have been derived elsewhere and we refer the user to full derivations (Elson and Magde 1974). Below we have the accepted definition of the 3-dimensional diffusion correlation function G(j,t):

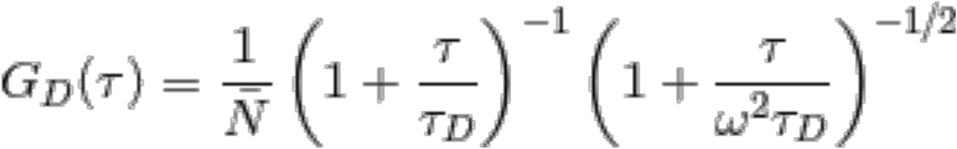

with parameters τ_D_(the characteristic transit time of the observed molecules through the observation spot), Ñ the number of particles, ω ^2^ a constant for connecting the transit time in 3-dimensions to 2-dimensions. FoCuS-scan has a variety of extensions to the classical diffusion equation with options that can be customised to describe multiple diffusion species, triplet states and anomalous diffusion:

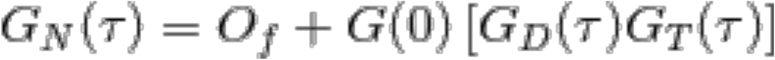

where Of is the offset, GN(0) is the amplitude of the correlation function (1/ Ñ, inverse average number of particles), GD is the diffusion model (2- or 3-dimensions) and GT is the optional triplet state. Triplet state equations are used to model the cases when the fluorophores under investigation have dark-states that are induced by the imaging regime (Widengren, Mets et al. 1995, Eggeling, Widengren et al. 1998). Within FoCuS-scan there is the facility to include one or more triplet states in the equation. Anomalous diffusion is a special situation where the diffusion regime deviates from the ideal relationship, for example, when studying diffusion using a super-resolution imaging regime. Further details for the specific components of this model are available in the FoCuS-scan supplementary manual.

#### Data visualization

Scanning FCS experiments typically produce large quantities of data and with this comes the requirement for powerful visualization tools to handle this data in an appropriate way. FoCuS-scan has multiple tools that allow users to analyse and dissect their data effectively including, filters, scatter plots and histogramming methods for visualizing populations of data. A detailed guide for these tools is available in the supplementary material section.

### Results

#### Scanning FCS simulation and live cell comparison

To validate FoCuS-scan and to understand the characteristics of diffusion across a range of physiologically and experimentally relevant rates we simulated Brownian motion and confocal scanning acquisition in 2-dimensions and compared the data to experiments performed on live cells under similar settings looking at the diffusion of a fluorescent DPPE-Atto647N lipid analogue (1,2-dipalmitoyl-sn-glycero-3-phosphoethanolamine tagged with the organic dye Atto647N) in the membrane. Figure 1 depicts experimentally acquired data from the DPPE-Atto647N lipid analogue in the plasma membrane of a Jurkat T cell (Fig. 1A) and a sample of the corresponding intensity and integrated time-series from an elliptical scan trajectory (Fig. 1B). Figure 1C shows the schematic for a scanning FCS simulation with very similar settings to the live cell experiment (including photon counting noise) and the corresponding sample intensity and integrated intensity time-series (Fig. 1D). Upon correlation of the live-cell or simulation data, correlation functions for each pixel are produced and can be presented as a plot of functions (Fig. 1E and 1G) or as a correlation carpet (Fig. 1F and 1H). For correlation carpets the maximum correlation are usually normalised to 1.0 along for each function so that the heterogeneity in the transit times can be easily observed. The distribution and form of the correlation functions are very similar between the actual live-cell data and also those generated from the simulation, and so it is possible to explore the fundamental phenomena of scanning FCS using these simulations.

Simulated intensity carpets for molecules diffusing in 2-dimensions (i.e. on membranes) with diffusion coefficients of D = 1.0, 0.5, 0.2 and 0.05 μm^2^/s were generated, respectively, with in each case 120 molecules being simulated for a duration of 30 s, a dwell time of 0.002 ms and a scan rate of 1800 Hz. 10 carpets were generated in total for each condition, resulting in 640 measurement points per different diffusion coefficient condition. Using a FCS-based analysis pipeline, each measurement point gave a value of the average transit time through the observation spot, and we could thus determine the distribution of transit times along with median values and variances (or standard deviations). From the simulated data it is clear that simulations for lower diffusion coefficients, i.e. slower diffusion (e.g. D = 0.05 μm^2^/s) exhibit larger median values of transit times and consequently a much greater absolute variance in values when compared to distributions generated from higher diffusion coefficients, i.e. faster diffusion (e.g. D = 1 μm^2^/s) (Fig. 2A).

**Figure 2:**
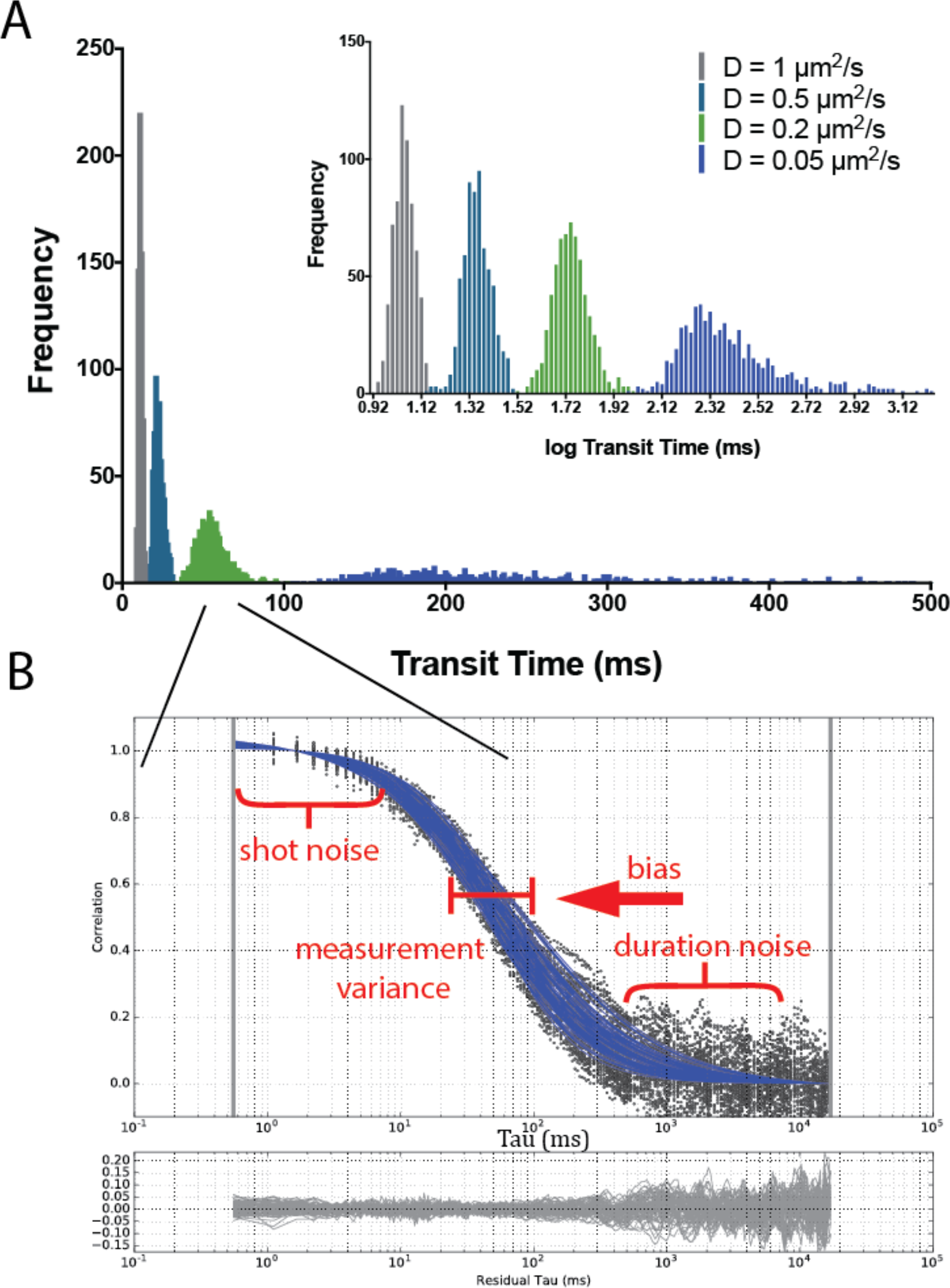
Simulated data generated across physiological ranges exhibit varying degrees of noise and statistical variance. **A**) Histogram of values of transit times determined via correlation analysis from 640 measurements from scanning FCS simulations with 1.0 μm^2^/s (**grey**), 0.5 μm^2^/s (**turquoise**), 0.2 μm^2^/s (**green**) and 0.05 μm^2^/s (**blue**) diffusion rates, representing 10x carpets from each simulation. (**Inset**) The same data but with natural logarithm of transit time values. **B**) Correlation functions from 0.2 μm^2^/s diffusion simulations (**black dots**) with parameterised fits (**blue lines**) with different forms of noise annotated on the different parts of the curve, and (**lower panel**) residuals from curve fitting (**grey lines**).

The broadening of the distribution of determined transit times is actually so dramatic for the slow-diffusion data that a logarithmic axis is required to adequately visualize the distribution across the whole range of simulated data (Fig 2A inset). Note that the simulations depicted in Figure 2A included no additional measurement noise in terms of shot noise, photon counting or photobleaching, yet, they still exhibit a high variance in terms of the observed transit values. This observation proved that in a typical scanning FCS paradigm powerful statistical effects are influencing the experimental output. Furthermore, in addition to the high degree of variance at the slower diffusion speeds (0.05 μm^2^/s) a systematic error (bias) is evident with the deviation of the population median deviating away from the theoretical value being simulated, as outlined further on. Figure 2A depicts a simulation of duration 30 s that is a realistic duration for an experiment performed on live cells and so a practical understanding of the observed variance phenomenon is very important.

Through using these simulations it is possible to accurately characterise the distributions and describe the affect that statistical variance and sampling will have on a likely experiment. As discussed in the introduction, certain phenomena will impact on different aspects of correlation and affect the statistics of the resultant function. Figure 2B shows a typical scanning FCS experiment with 64 curves generated from a single simulated carpet. Highlighted in this plot are the areas known to be affected by specific phenomena (e.g. shot/duration noise, bias, measurement variance). The top of each curve is susceptible to effects caused by poor signal-to-noise due to poor photon yield and other effects (annotated ‘shot noise’) (Koppel 1974). If noise is evident at the bottom of the curve, this is due to an insufficient experimental duration compared to the measured transit time introducing variance (Schätzel, Drewel et al. 1988, Saffarian and Elson 2003) and is annotated as ‘duration noise’. Variance in the middle of curve, in terms of the positions of the curve inflexion points (or transit times), is a consequence of non-convergence of the underlying molecules being measured, and again is dependent on the duration of the experiment and the speed of the molecule being measured (‘measurement variance’). A systematic shift (‘bias’) is also a consequence of the duration of an experiment, but also of the correlation methodology as a whole, as we see later, measurements made using single molecule tracking do not suffer this bias (Schätzel, Drewel et al. 1988). Having researched the possible causes of variance in our data it was clear that our data could be affected by any number of them, so we sort to delineate each influence for the benefit of our own understanding and that of subsequent FoCuS-scan users.

#### Establishing the cause of the variance and systematic error in the transit time distributions

To test whether the observed variance and systematic errors were a product of correlation, or intrinsic to the system being studied, single molecule tracking was applied to track molecules entering the observation volume for each simulation (Fig. 3). Rather than the FCS approach of integrating the particles with the PSF, the time for each particle was determined for the duration each particle was in the observation volume (1/e^2^ radius, 212.33 nm, as threshold), and the distance between the exit and entry point measured also for each event (Fig 3A). Given the duration and effective distance of each particle entering the observation volume it was straightforward to calculate the transit time of each particle and the average time for the entire ensemble. Repeating this for each simulation measurement, generated a population of measurements very similar to those generated through the correlation method. Figure 3B compares the distribution of values of transit times generated from simulated diffusion (D = 0.2 μm^2^/s) and analyzed by correlation (green) and single molecule tracking (black). The distributions obtained from the single molecule tracking analysis revealed strong differences to those obtained from the correlation analysis, especially differences in their median value and their overall shape. These differences and their comparison to the theoretical values used in the simulations allowed us to highlight systematic and convergence artifacts associated with the determination of transit times via (scanning) FCS.

**Figure 3:**
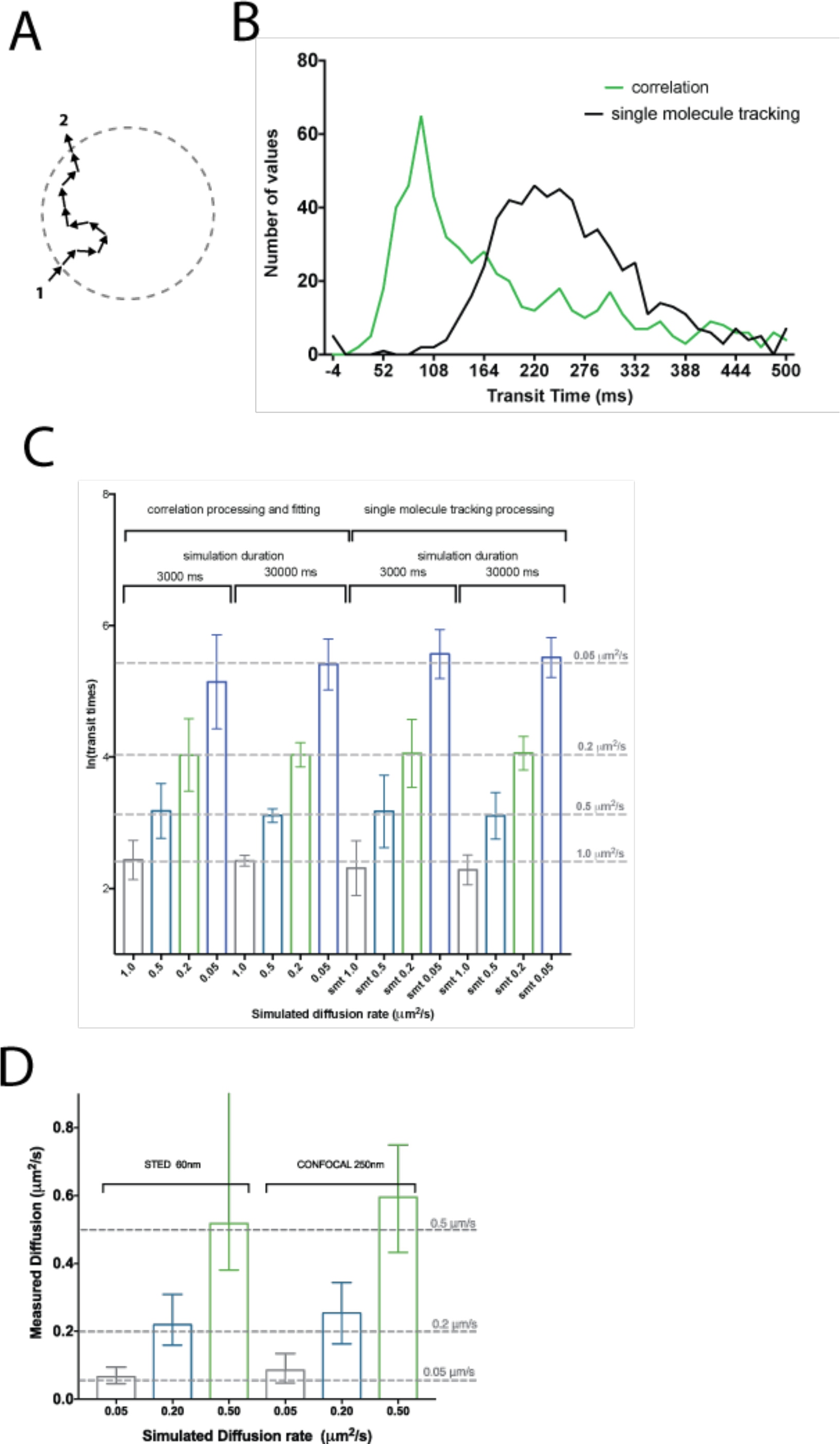
Variance in transit time measurements is underlying statistical variance of particle movement, bias is characteristic of correlation methodology. Comparison of transit time populations from different analysis methods and for different measurement durations of the simulations. A) Schematic representation of the single molecule tracking analysis method. The Euclidean distance between where the particle entered (1) and exited (2) the observation spot area (circle marking the observation spot/PSF boundary, 1/e^2^ diameter of the 250 nm FWHM PSF as border) is measured and divided by the duration. B) Histogram (bin size=14, D=0.05 μm^2^/s) of transit time values from 640 simulated measurements determined with single molecule tracking method (black) and with correlation analysis (green). C) Median and standard deviations (error bars) of transit time values from 640 simulated measurements with different theoretical simulated diffusion coefficients and durations and determined with the different analysis methods (single molecule tracking and correlation), as labelled. D) Median values of diffusion coefficients determined from 640 simulated measurements with a spot size of 60 nm FWHM (STED) compared to a spot size of 250 nm FWHM (confocal) and different diffusion coefficients of 1.0 μm^2^/s (grey), 0.5 μm^2^/s (turquoise), 0.2 μm^2^/s (green) and 0.05 μm^2^/s (blue). Error bars represent inter-quartile range of obtained values. Grey dashed lines in C and D represent theoretically expected values, and deviation of bars from nearest line represents bias.

#### Explanation for the systematic and convergence artifacts in the transit time distributions

When modeling stochastic random processes over limited time periods there will always be some deviation in each modeled measurement from those acquired over an infinite time duration. For scanning FCS this deviation is clearly evident as measurement variance, this is because of the limited acquisition time with respect to the relatively slow diffusion and thus rate of molecular species being investigated. As highlighted, through simulating single molecule tracking we have been able to circumvent the possibility that the correlation processing and fitting is influencing the broadening of the measured transit times. A closer inspection of the median values and variances of the transit times generated from either analysis method (FCS and single molecule tracking) for data simulated for differently moving molecules (D = 0.05 – 1.0 μm^2^/s) and different total acquisition times (3 and 30 s) revealed a high degree of similarity (Fig. 3C), despite the general difference in the original distribution (Fig. 3B). Both FCS and single molecule tracking have a characteristic precision when looking at a population of measurements, which decreases with total acquisition time. For example, for data simulated with D = 0.05 μm^2^/s FCS analysis yielded median values (and standard deviations) of transit times of 161 ± 175 ms after 3000 ms and 212 ± 116 ms after 30000 ms simulated duration, and single molecule tracking analysis 253 ± 127 ms after 3000 ms and 246 ± 92 ms after 30000 ms simulated duration. Consequently, both methods missed some accuracy to reproduce the true input transit time of 225.4 ms, especially for short acquisition times. Note that this inaccuracy is less a problem for fast moving molecules such as for the data simulated with D = 0.5 and 0.2 μm^2^/s.

This inaccuracy shows that there is an inherent problem of measuring the kinetics of molecules crossing the detection volume. If molecules are diffusing across an area, some will take longer times than others, due to statistical variance. If data is acquired for a short period of time only it is likely that one may not probe enough transits to find the true average, as by chance one may have observed only a limited set of events, that happened to be on average quicker or slower than the true average generated from an infinite measurement time. The longer one monitors, the more accurate each individual assessment of the average will be and therefore the tighter the resulting population of measurements will become. This convergence phenomenon is dependent on the speed of the particle under study, the length of the study and also the size of the area used. As pointed out (Fig. 3C), in the simulations the slower moving particles result in wider distributions, and the longer the duration of the measurement, the tighter the distribution becomes, no matter of the speed. Corrections developed in the past will only tackle the noise across the correlation function, also caused by the lack of recording duration (Wohland, Rigler et al. 2001, Saffarian and Elson 2003), rather than adjusting the underlying population of transit times to point at which they converge. While dependencies of the noise of correlation data on the measurement time and general diffusion speed have been indicated before (Koppel 1974, Saffarian and Elson 2003), to our knowledge this is the first time that simulations have been run at sufficient numbers and at slow enough diffusion and thus molecular rates to visualize population transit time variance of this scale, though this relationship has been observed in experimental measurements (Kask, Günther et al. 1997). Through understanding the underlying cause of the variance it is possible to devise strategies to cancel it out.

#### Systematic error associated with the correlation method

Due to the calculation of the correlation function, the FCS data, in addition to the basic variance pointed out in the previous paragraph, also suffer from systematic deviations when the experimental duration is limited, reducing the accuracy of the FCS-based analysis further. This phenomena has been previously described as correlation ‘bias’ (Schätzel, Drewel et al. 1988) and has been pointed out in previous high-throughput screening experiments (Kask, Günther et al. 1997, Eggeling, Brand et al. 2003). This ‘bias’ is highlighted by our current observations that distributions of transit time values generated from single molecule tracking analysis are more accurately describing the true theoretical average (as input by the simulations) than those generated from FCS (Fig. 3C).

Let us again consider the example of data simulated for D = 0.05 μm^2^/s, with a true theoretical transit time of 225.4 ms. The FCS analysis of this data yields a median transit time and standard deviation of 161 ± 175 ms after 3000 ms duration simulation showing poor accuracy but achieves improved accuracy with a value of 212 ± 116 ms after 30000 ms. The same comparison for the analysis of this data set using single molecule tracking analysis achieves 253 ± 127 ms and 246 ± 92 ms for 3000 and 30000 ms simulation time, respectively, showing almost no change in median value and a smaller increase in standard deviation with reduced measurement duration. Note that this prominent difference between FCS and single molecule tracking analysis in decreased accuracy for shorter measurement times diminishes for the simulations with faster moving molecules (e.g. D = 0.5 – 1 μm^2^/s, Fig. 3C). In summary, our simulations highlight that for short measurement times on especially slow moving particles, the correlation bias is more of a problem than the variance, and some corrective strategies are required to reduce the bias.

#### Removing systematic bias and improving convergence

There are several potential solutions for reducing the bias and for assisting in the convergence of the measurements. The first and most obvious is to increase the measurement time for a longer more stable measurement. The drawback of this approach is that biological specimens such as cells may become damaged due to phototoxic effects, might move during acquisition, or the dynamics under-study may change, and overall this limits the number of measurements that can be made. The second is to apply a correction to the correlation function which is possible through the calculation and application of a 1st order bias factor (Saffarian and Elson 2003). The third proposed technique is to reduce the size of the observation volume, which can be achieved by applying FCS on a super-resolution STED microscope (Kastrup, Blom et al. 2005, Eggeling, Ringemann et al. 2009, Rossow, Sasaki et al. 2010, Honigmann, Mueller et al. 2014, Waithe, Clausen et al. 2015). Reducing the observation spot size of the acquisition improves the accuracy and precision of scanning FCS measurements. In Figure 3D the effects of reducing the observation volume through using a STED microscope are simulated on datasets only 3000 ms in length. Clear gains in accuracy are seen for slower transit times when compared to analysis using conventionally sized confocal observation volumes. For simulated experiments with molecules diffusing with D = 0.05 and 0.2 μm^2^/s we using the FCS analysis obtained median values of diffusion coefficients of 0.066 and 0.22 μm^2^/s, respectively, for an observation spot of 60 nm in diameter (STED) and 0.086 and 0.25 μm^2^/s, respectively, for the conventional confocal observation spot (250 nm in diameter). The use of the 60 nm large observation spots is however not advisable for very fast moving molecules with diffusion coefficients of 0.5 μm^2^/s or greater because in these cases the transit times are too close to the temporal resolution of the scan line (in this case 1800 Hz, i.e. 0.556 ms between line repetitions); however a higher scan rate would correct this (e.g. by using a fast resonance scanner). For slower diffusing species (D = 0.2-0.05 μm^2^/s), reducing the observation spot size as in the presented case can be equivalent to a 10 x increase in experiment duration, making it possible to take accurate unbiased measurements in a fraction of the time.

#### Noisy data, error metrics and data visualization

As discussed in the introduction, several studies have investigated the impact of noise in terms of assessing the quality of correlation functions. One of the standard approaches is to establish the signal-to-noise ratio for a given correlation function S/N = G(t)/var(G(t))^0.5^ (Koppel 1974). When we applied a normalized variant of the signal-to-noise formula (Krichevsky and Bonnet 2002) to our data taken for 30000 ms, we found that the average S/N for our measurements was linearly dependent on the measured transit time (Fig 5A). This means that the longer the transit time the less certain one is as to the true value of that data-point as well as explaining why the distributions are log-normal in appearance. Each calculated point is effectively convolved with an error that increases with transit time producing non-symmetrical distributions (Can and Log 2001). This effect is very likely linked again to the length of the measurement duration and the effect this can have on the variance of a function due to slower moving particles residing longer in the detection volume. Histogramming techniques, although reliable for binning and visualization of data, assume in most cases a finite bin size. Because the distributions we observe can span large transit time margins we wanted a method of binning and visualization which could take into account the increased margin for uncertainty with longer transit times.

#### Density kernel estimation bandwidth calculations

Density kernel estimation is an alternate technique to histogramming used for visualizing and also for estimating the probability density distribution of a random variable. With this method each data point is represented with a kernel (typically a Gaussian) and the bandwidth of the kernel is set to a static width for each point or is varied depending on a suitable metric. Because the kernel size can be set dynamically, it is advantageous over conventional histogram methods, which are restricted to a static bin and cannot be tuned to individual data points. Here we trialed using density kernel estimation with a variable bandwidth proportional to the level of error we obtained from three different error metrics. Each transit time data point was represented as a Gaussian kernel of standard deviation equal to the error associated with the calculation of that particular transit time measurement. In addition to the signal-to-noise estimation we implemented two additional metrics for calculating the variance of our transit time data populations and first wanted to establish there suitability as error metrics for correlation. The first method was to use the standard deviation calculated for our transit time parameter during curve fitting using the lmfit python library. This method was very efficient to calculate as this error parameter accompanies each of our fitting operations. Our second method was to perform bootstrapping on our correlation data, followed by calculating the standard deviation of the transit times generated across the bootstraps. Bootstrapping is a commonly used statistical test for assessing the accuracy of an estimator, in this case the estimator being the correlation function. Bootstrapping is often used when parametric inference is impossible or difficult, as is the case when calculating correlation factor error. In this method we take each of our correlated measurements and then randomly sample with replacement the function to form a bootstrap sample. This process is repeated a number of times (we use 5-100x) and the transit time calculated for each of the distributions, and the standard deviation calculated across the population. We found that for both the methods trialed, the curve fitting standard deviation parameter and the bootstrapping method, both gave error measures consistent with the signal-to-noise statistic calculated for each curve (Fig. 4B-C). Each of these methods had a positive linear relationship with regard to the transit time for each of the generated datasets and a distribution similar in form to the signal-to-noise estimation and so we deemed them good potential candidates to express the error in our density kernel visualization.

**Figure 4:**
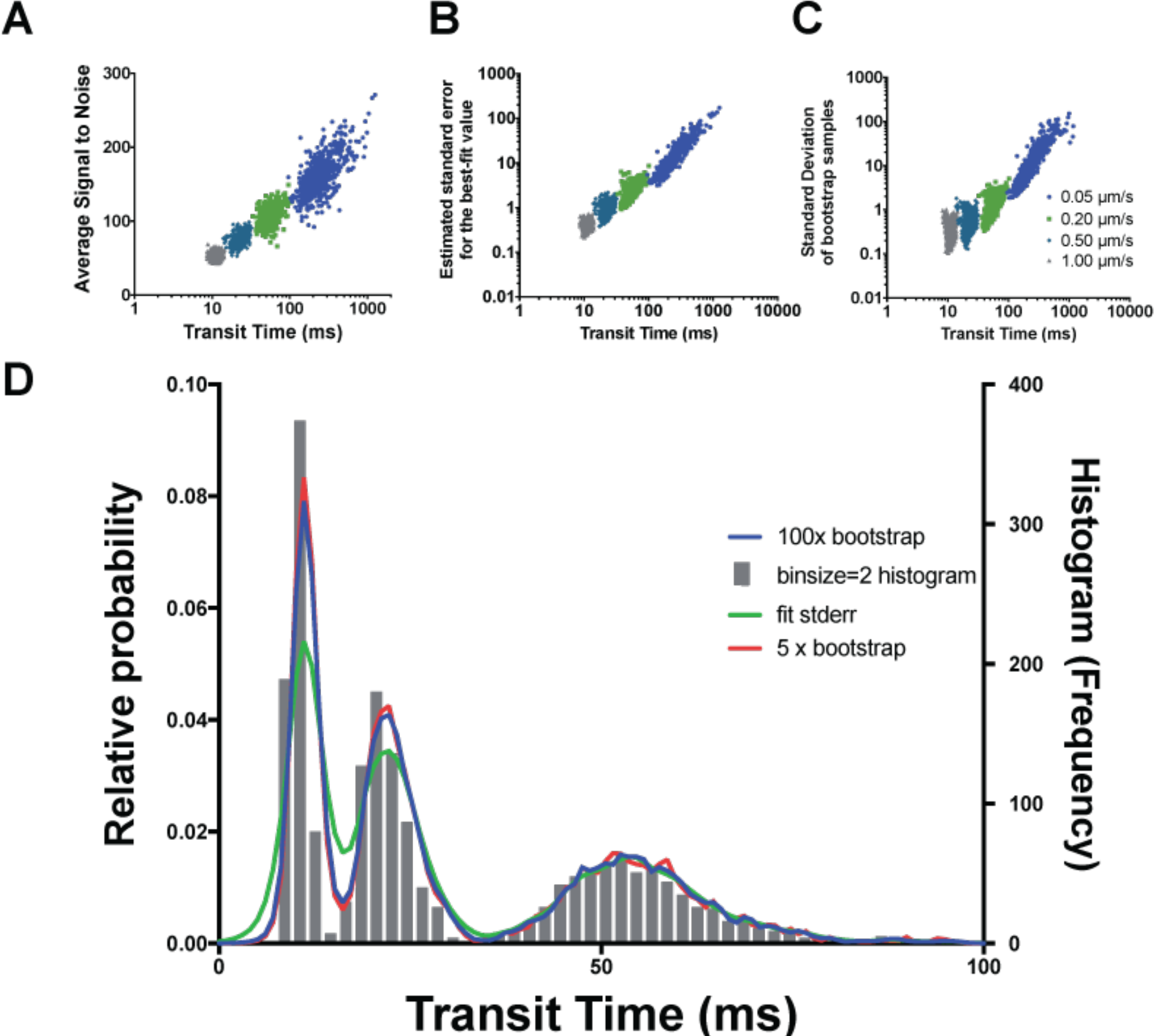
Density kernel estimation enables visualization of transit time distributions across scales. **A-C**) Scatter plots comparing transit time measurements from 640 simulations (30 s duration, 120 mol.) with diffusion coefficients of 1.0 μm^2^/s (**grey**), 0.5 μm^2^/s (**turquoise**), 0.2 μm^2^/s (**green**) and 0.05 μm^2^/s (**blue**) with different error metrics, signal-to-noise (**A**), correlation fit standard deviation (**B**) and standard deviation of 100x bootstrap samples (**C**). **D**) Density kernel estimation of the data of A (shown here as histogram, with binsize=2) with different error metrics compared to Histogram visualization method with widths corresponding to 100x bootstrap (**blue-line**), 5x bootstrap (**red-line**), correlation fit standard deviation (**green-line**).

#### Density kernel estimation visualization

Once we have established our potential error metrics we trialed each on the data. To compare our methods we first amalgamated data into a single transit time distribution from simulations of 1.0 μm^2^/s, 0.5 μm^2^/s and 0.2 μm^2^/s diffusion rates. Next we histogrammed the data using a bin-size = 2, optimized for the transit time relating to 1.0 μm^2^/s. The average signal-to-noise ratio data produced kernels of a width which was far too large (data not shown), however the curve fit error and the bootstrap methods both produced density kernel visualizations which were highly similar to the input data with the histogram (optimized for 1.0 μm^2^/s) (Fig. 4D). We found that the bootstrap method produced a distribution remarkably similar to the histogram distribution and was tighter than the curve fit error method. We also show that as little as 5x bootstraps is necessary to accurately represent the distribution of the data, as both this distributions are highly similar. Density Kernel estimation is advantageous as the output function can be interpolated to any degree of accuracy and also there is no requirement to define a bin-size, making it advantageous over histogramming. In summary, the bootstrapping method, as an error metric, closely matches the trends of more well defined metrics of variance such as signal-to-noise and can be used as an appropriate scale invariant metric for density kernel bandwidth estimation, which is a superior visualization method when compared to histograms.

#### Photobleaching correction

Due to the repetitive nature of FCS measurements, some photobleaching is likely to occur during the course of a scanning FCS acquisition experiment. Scanning FCS suffers from photobleaching less than point FCS due to the sequential nature of the line scanning and therefore reduced continuous exposure of any one point to the laser,(Donnert, Eggeling et al. 2009) however some photobleaching can still occur. Photobleaching represents loss of information and corrections should be applied with care. Fortunately, due to the differential nature of photobleaching, in that it affects immobile fluorescent fractions much more than mobile fractions, it is possible to extract the signal from the mobile fraction near perfectly despite a seemingly large effect on the unwanted immobile fraction. The main perturbation which occurs to a correlation signal as a consequence of photobleaching is due to the effect that photobleaching has on the lower temporal frequencies of the correlation function and the overall impact this has on the normalisation of the correlation function (Icenogle and Elson 1983, Hess and Webb 2002). The photobleaching occurs on a different time-scale to the mobile fraction which are left relatively unaffected except that the normalisation of the correlation function as whole is perturbed which can affect the fitting of the correlation function. This effect is clearly visualised in Figure 5. In Fig. 5A there is a simulated intensity time-series of diffusing molecules with no photobleaching and in Fig. 5B the same intensity time-series is depicted but with the addition of photobleaching simulated through the addition of a mono-exponentially decaying function. When these signals are correlated we obtain correlation functions depicted in figure 5C with the correlation curves generated from the non-photobleaching signal (green) and the photobleached signal (black). The correlation curve generated from the photobleaching signal lacks an inflexion point due to the perturbation that the photobleaching has on the normalisation of the signal. Through using either of the two photobleaching correction algorithms it is possible to restore the curve to something much more similar to the non-photobleached original (Fig. 5C, red and blue lines). Correction method 1 involves fitting a mono-exponential function to the intensity signal of the whole intensity carpet and then a correction is applied to the intensity signal for the intensity over time (see methods for more details) (Ries, Chiantia et al. 2009). This has the effect that the normalisation of the signal is corrected directly but will introduce artefacts if the photobleaching signal is not mono-exponential or differs between positions of the carpet. Correction method 2 (local averaging) involves breaking up the intensity signal for each pixel up into time-series of shorter length, correlating each section and then averaging the output function (Wachsmuth, Conrad et al. 2015). Although the time-series sequence is reduced in length, which will increase the variance of the species being studied, the averaging of the correlated sequences will reduce the broadening effect this has on the population data. However, when used for slower diffusing species it is likely that some bias may be introduced as a result of the temporally shortened sequence. This approach works well however for faster moving species as the photobleaching affects on the lower temporal frequencies will be filtered out through using sections of shorter length. Analysis of a population of data as shown in Figure 5D shows that, in terms of restoring the signal, so that the transit time measurements are accurate, both correction method 1 and 2 are far superior to no correction, and correction method 2 produces a distribution most similar to the original non-photobleached population with the closest median population transit time. In summary although photobleaching should be avoided initially through design of the experiment (suitable fluorophores, low laser power etc.), photobleaching correction methods are highly effective at reducing the impact that the photobleaching has on the data distribution. FoCuS-scan has two accepted methods that are effective at correcting likely photobleaching artefacts and these should be used when appropriate to restore a signal through correction of its normalisation.

**Figure 5:**
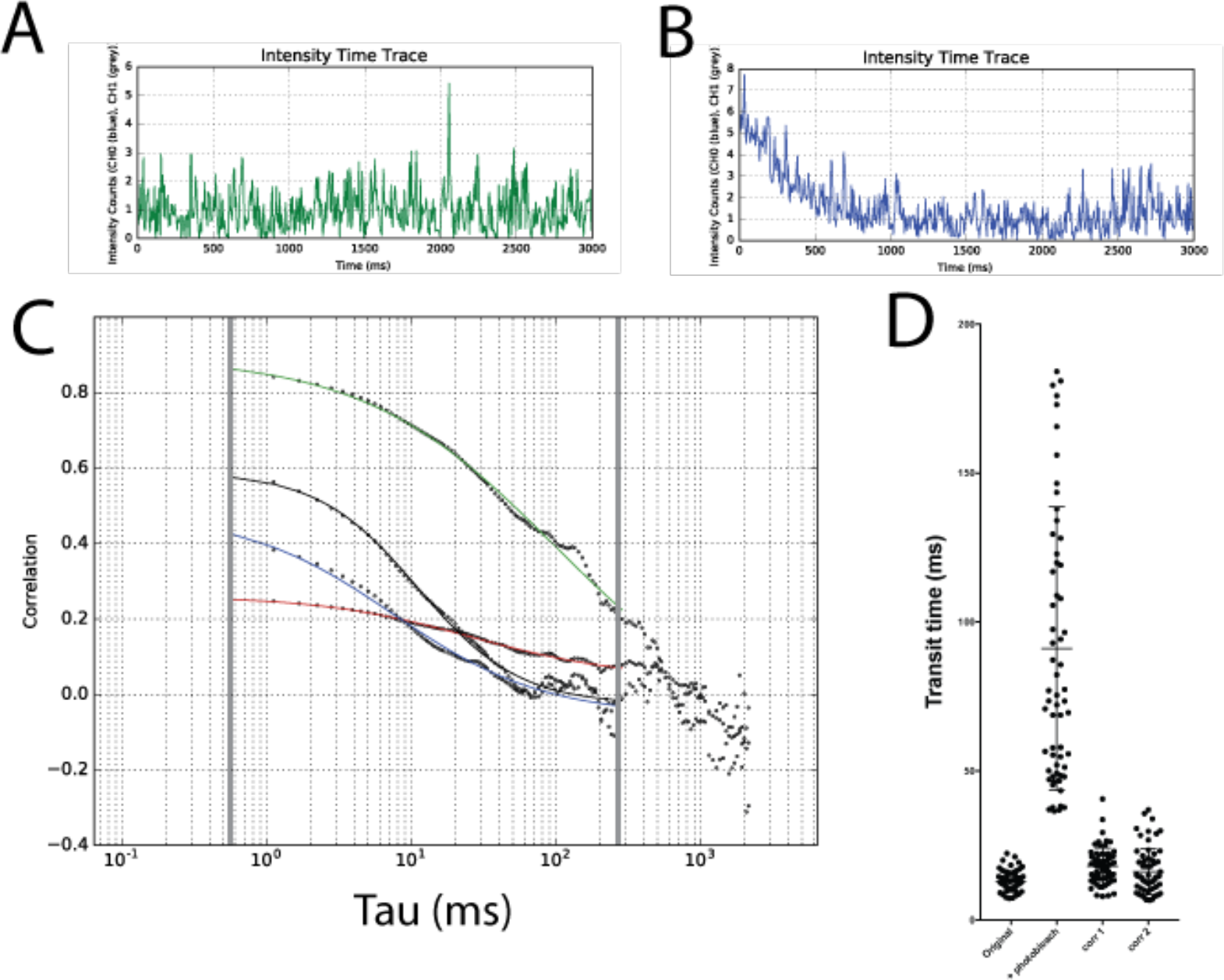
Photobleaching dramatically affects transit time measurements but can be restored through suitable corrections. **A,B**) Intensity time-series from a simulated experiment (D = 0.5 μm^2^/s, duration 3000 ms), integrated over all pixels from simulation without (**A**) and with photobleaching (**B**). (**C**) Respective correlation curves from the simulation without (**black line**) and with photobleaching (**green line**, analyzed in the conventional way), and in the latter case with correction method 1 (**red line**) and correction method 2 (**blue line**). **D**) Whisker plots of values of transit times determined from 64 simulated measurements generated with and without photobleaching, original data and analysis without correction and with correction methods 1 and 2, as labeled.

#### Conclusion

Scanning FCS is a powerful technique for establishing the spatio-temporal kinetics of diffusion in live cells or potentially in supported lipid bilayers. So far the technique has only seen limited use within the life sciences due to the requirements of specialist equipment and analysis software as well as the need for validation within this area. With FoCuS-scan it is now possible to process and analyse data acquired on conventional confocal scanning microscopes. Furthermore, scanning FCS has great potential to be applied in future to also quantify more complex diffusion and reaction dynamics of molecules such as molecule oligomerisation and/or transient binding events in living cells. Such analysis will require additional strategies to fulfil the obvious demand in cell-biology to systematically and statistically robustly quantify these processes. This article describes the performance of scanning FCS across a broad range of biological relevant diffusion length scales and highlights the limitations and practical considerations when performing these experiments.

## Acknowledgements

We acknowledge the WIMM for infrastructure support. We thank the Wolfson Imaging Centre Oxford (especially Dr. Christoffer Lagerholm) and the MICRON Advanced Bioimaging Unit Oxford (supported by the Wellcome Trust Strategic Award 091911) for access to equipment and assistance with data acquisition and analysis. The work was supported by the Wolfson Foundation, the Medical Research Council (MRC, grant number MC UU 12010/unit programmes G0902418 and MC UU 12025), MRC/BBSRC/EPSRC (grant number MR/K01577X/1), Wellcome Trust (Strategic Award 091911 and grant ref 104924/14/Z/14), Deutsche Forschungsgemeinschaft (Research unit 1905), and internal University of Oxford funding (EPA Cephalosporin Fund and John Fell Fund). JBdlS acknowledges his support by a Marie Curie Career Integration Grant, NanodynacTCELLvation (grant number PCIG13-GA-2013-618914). We thank Dr. Marco Fritzsche and Dr. Silvia Galiani for their proofreading and advice whilst editing.

## Bibliography

Brinkmeier, M. , K. Dörre , J. Stephan and M. Eigen (1999). “Two-beam cross-correlation: a method to characterize transport phenomena in micrometer-sized structures.” Analytical chemistry 71(3): 609–616.

Can, W. and O. Log (2001). “Log-normal distributions across the sciences: Keys and clues.” BioScience 51(5).

Colyer, R. A. , G. Scalia , T. Kim , I. Rech , D. Resnati , S. Marangoni , M. Ghioni , S. Cova , S. Weiss and X. Michalet (2010). High-throughput multispot single-molecule spectroscopy. BiOS, International Society for Optics and Photonics.

de la Serna, J. B. , G. J. Schütz , C. Eggeling and M. Cebecauer (2016). “There Is No Simple Model of the Plasma Membrane Organization.” Frontiers in cell and developmental biology 4.

Digman, M. A. , P. Sengupta , P. W. Wiseman , C. M. Brown , A. R. Horwitz and E. Gratton (2005). “Fluctuation correlation spectroscopy with a laser-scanning microscope: exploiting the hidden time structure.” Biophysical journal 88(5): L33–L36.

Donnert, G. , C. Eggeling and S. W. Hell (2009). “Triplet-relaxation microscopy with bunched pulsed excitation.” Photochemical & Photobiological Sciences 8(4): 481–485.

Eggeling, C. , L. Brand , D. Ullmann and S. Jäger (2003). “Highly sensitive fluorescence detection technology currently available for HTS.” Drug discovery today 8(14): 632–641.

Eggeling, C. , C. Ringemann , R. Medda , G. Schwarzmann , K. Sandhoff , S. Polyakova , N. Belov , B. Hein , C. von Middendorff and A. Schönle (2009). “Direct observation of the nanoscale dynamics of membrane lipids in a living cell.” Nature 457(7233): 1159–1162.

Eggeling, C. , J. Widengren , R. Rigler and C. Seidel (1998). “Photobleaching of fluorescent dyes under conditions used for single-molecule detection: Evidence of two-step photolysis.” Analytical chemistry 70(13): 2651–2659.

Ehrenberg, M. and R. Rigler (1974). “Rotational brownian motion and fluorescence intensify fluctuations.” Chemical Physics 4(3): 390–401.

Elson, E. L. and D. Magde (1974). “Fluorescence correlation spectroscopy. I. Conceptual basis and theory.” Biopolymers 13(1): 1–27.

Fahey, P. , D. Koppel , L. Barak , D. Wolf , E. Elson and W. Webb (1977). “Lateral diffusion in planar lipid bilayers.” Science 195(4275): 305–306.

Galiani, S. , D. Waithe , K. Reglinski , L. D. Cruz-Zaragoza , E. Garcia , M. P. Clausen , W. Schliebs , R. Erdmann and C. Eggeling (2016). “Super-resolution Microscopy Reveals Compartmentalization of Peroxisomal Membrane Proteins.” Journal of Biological Chemistry 291(33): 16948–16962.

Hebert, B. , S. Costantino and P. W. Wiseman (2005). “Spatiotemporal image correlation spectroscopy (STICS) theory, verification, and application to protein velocity mapping in living CHO cells.” Biophysical journal 88(5): 3601–3614.

Hess, S. T. and W. W. Webb (2002). “Focal volume optics and experimental artifacts in confocal fluorescence correlation spectroscopy.” Biophysical journal 83(4): 2300–2317.

Honigmann, A. , V. Mueller , H. Ta , A. Schoenle , E. Sezgin , S. W. Hell and C. Eggeling (2014). “Scanning STED-FCS reveals spatiotemporal heterogeneity of lipid interaction in the plasma membrane of living cells.” Nature communications 5.

Icenogle, R. and E. Elson (1983). “Fluorescence correlation spectroscopy and photobleaching recovery of multiple binding reactions. II. FPR and FCS measurements at low and high DNA concentrations.” Biopolymers 22(8): 1949–1966.

Kask, P. , R. Günther and P. Axhausen (1997). “Statistical accuracy in fluorescence fluctuation experiments.” European biophysics journal 25(3): 163–169.

Kastrup, L. , H. Blom , C. Eggeling and S. W. Hell (2005). “Fluorescence fluctuation spectroscopy in subdiffraction focal volumes.” Physical review letters 94(17): 178104.

Koppel, D. E. (1974). “Statistical accuracy in fluorescence correlation spectroscopy.” Physical Review A 10(6): 1938.

Krichevsky, O. and G. Bonnet (2002). “Fluorescence correlation spectroscopy: the technique and its applications.” Reports on Progress in Physics 65(2): 251.

Lingwood, D. and K. Simons (2010). “Lipid rafts as a membrane-organizing principle.” science 327(5961): 46–50.

Magde, D. , E. Elson and W. W. Webb (1972). “Thermodynamic fluctuations in a reacting system—measurement by fluorescence correlation spectroscopy.” Physical Review Letters 29(11): 705.

Müller, P. (2012). “Python multiple-tau algorithm.”

Müller, P. , P. Schwille and T. Weidemann (2014). “Scanning fluorescence correlation spectroscopy (SFCS) with a scan path perpendicular to the membrane plane.” Fluorescence Spectroscopy and Microscopy: Methods and Protocols: 635–651.

Newville, M. , A. Nelson , T. Stensitzki , A. Ingargiola , D. Allan , Y. Ram , C. Deil , G. Pasquevich , T. Spillane and P. A. Brodtkorb (2016). “LMFIT: non-linear least-square minimization and curve-fitting for Python.” Astrophysics Source Code Library.

Papadopoulos, D. K. , A. J. Krmpot , S. N. Nikolić , R. Krautz , L. Terenius , P. Tomancak , R. Rigler , W. J. Gehring and V. Vukojević (2015). “Probing the kinetic landscape of Hox transcription factor–DNA binding in live cells by massively parallel Fluorescence Correlation Spectroscopy.” Mechanisms of development 138: 218–225.

Petersen, N. O. , P. L. Höddelius , P. W. Wiseman , O. Seger and K. Magnusson (1993). “Quantitation of membrane receptor distributions by image correlation spectroscopy: concept and application.” Biophysical journal 65(3): 1135–1146.

Ries, J. , S. Chiantia and P. Schwille (2009). “Accurate determination of membrane dynamics with line-scan FCS.” Biophysical journal 96(5): 1999–2008.

Rigler, R. and J. Widengren (1990). “Ultrasensitive detection of single molecules by fluorescence correlation spectroscopy.”

Rossow, M. J. , J. M. Sasaki , M. A. Digman and E. Gratton (2010). “Raster image correlation spectroscopy in live cells.” Nat. Protocols 5(11): 1761–1774.

Ruan, Q. , M. A. Cheng , M. Levi , E. Gratton and W. W. Mantulin (2004). “Spatial-temporal studies of membrane dynamics: scanning fluorescence correlation spectroscopy (SFCS).” Biophysical journal 87(2): 1260–1267.

Saffarian, S. and E. L. Elson (2003). “Statistical analysis of fluorescence correlation spectroscopy: the standard deviation and bias.” Biophysical journal 84(3): 2030–2042.

Sankaran, J. , M. Manna , L. Guo , R. Kraut and T. Wohland (2009). “Diffusion, transport, and cell membrane organization investigated by imaging fluorescence cross-correlation spectroscopy.” Biophysical journal 97(9): 2630–2639.

Schätzel, K. (1985). New concepts in correlator design. Inst. Phys. Conf. Ser.

Schätzel, K. , M. Drewel and S. Stimac (1988). “Photon correlation measurements at large lag times: improving statistical accuracy.” Journal of Modern Optics 35(4): 711–718.

Schwille, P. , J. Korlach and W. W. Webb (1999). “Fluorescence correlation spectroscopy with single - molecule sensitivity on cell and model membranes.” Cytometry 36(3): 176–182.

Sezgin, E. , I. Levental , S. Mayor and C. Eggeling (2017). “The mystery of membrane organization: composition, regulation and roles of lipid rafts.” Nature Reviews Molecular Cell Biology.

Wachsmuth, M. , C. Conrad , J. Bulkescher , B. Koch , R. Mahen , M. Isokane , R. Pepperkok and J. Ellenberg (2015). “High-throughput fluorescence correlation spectroscopy enables analysis of proteome dynamics in living cells.” Nature biotechnology 33(4): 384–389.

Wachsmuth, M. , T. Weidemann , G. Müller , U. W. Hoffmann-Rohrer , T. A. Knoch , Waldeck and J. Langowski (2003). “Analyzing intracellular binding and diffusion with continuous fluorescence photobleaching.” Biophysical journal 84(5): 3353–3363.

Waithe, D. , M. P. Clausen , E. Sezgin and C. Eggeling (2015). “FoCuS-point: software for STED fluorescence correlation and time-gated single photon counting.” Bioinformatics 32(6): 958–960.

Widengren, J. , U. Mets and R. Rigler (1995). “Fluorescence correlation spectroscopy of triplet states in solution: a theoretical and experimental study.” The Journal of Physical Chemistry 99(36): 13368–13379.

Widengren, J. and R. Rigler (1998). “Fluorescence correlation spectroscopy as a tool to investigate chemical reactions in solutions and on cell surfaces.” Cellular and molecular biology (Noisy-le-Grand, France) 44(5): 857–879.

Wohland, T. , R. Rigler and H. Vogel (2001). “The standard deviation in fluorescence correlation spectroscopy.” Biophysical journal 80(6): 2987–2999.

